# Diversity and disease: evidence for the monoculture effect beyond agricultural systems

**DOI:** 10.1101/668228

**Authors:** Alice K.E. Ekroth, Charlotte Rafaluk-Mohr, Kayla C. King

**Author notes:** **Authorship** A.K.E.E. and K.C.K conceived and designed the study. A.K.E.E. gathered the data and performed the statistical analysis with C.R-M. A.K.E.E. and K.C.K. wrote the paper. The authors have no competing interests.

## Abstract

Human activities are greatly reducing the genetic diversity of species worldwide. Given the prediction that parasites better exploit less diverse host populations, many species could be vulnerable to disease outbreaks. However, the widespread nature of the ‘monoculture effect’ remains unclear outside agricultural systems. We conducted a meta-analysis of 22 studies, obtaining a total of 66 effect sizes, to directly test the biological conditions under which host genetic diversity limits infectious disease in populations. Overall, we found broad support for the monoculture effect across host and parasite species. The effect was independent of host range, host reproduction, parasite diversity, and the method by which the monoculture effect was recorded. Conversely, we found that parasite functional group, virulence, and empirical environment matters. Together, these results highlight the general susceptibility of genetically homogenous populations to infection. Consequently, this phenomenon could become increasingly common and alarming for at-risk populations due to human-driven declines in genetic diversity and shifts in parasite distributions.

## Introduction

Most natural populations are genetically diverse (1). In host populations, genetic diversity is thought to increase the chance that one or more individuals in a population is resistant to infection, and thereby reduces the likelihood of a parasite encountering a susceptible host (2). Genetically homogenous host populations are conversely believed to be more vulnerable to infection given the uniformity of host susceptibility. This relationship between low genetic diversity and high disease incidence is referred to as the ‘monoculture effect’ (3).

The study of the monoculture effect in agricultural settings is extensive (4–6). A recent meta-analysis showed that with increased diversity in intraspecific cultivar mixtures disease presence is reduced and crop yields increased (6). However, we know little of the extent to which the monoculture effect can occur across species and environments in natural systems and beyond agricultural contexts. Crop plants are under artificial selection for high yield, and may therefore exhibit less genetic polymorphism than those in the wild.

Threats to genetic diversity are on the rise. Habitat alterations, pollution, and global temperature changes, as well as the restriction of species geographical ranges may lead to higher chances of genetic drift and reduced population genetic diversity (7). Consequently, populations might suffer diminished evolutionary potential (8) and increased inbreeding depression (9,10). Knowing whether there is an additional, and perhaps more immediate and intense, threat of outbreaks in these populations is crucial for disease management and species conservation approaches.

Theory has illuminated the dynamics of parasite spread (3,11–14) in diverse host populations as well as examined the level of diversity required to stop transmission (15,16). However, the generality of the monoculture effect in nature remains unclear for several reasons. Firstly, given the infection rates of some parasites can be determined by host density (2), the relative effects of density versus host genetic diversity need to be elucidated (16). Shrinking habitats, for example, can result in higher population densities (and lower resource availability) where parasites can transmit better due to more contact between hosts (17,18). Secondly, even when focusing on host genetic diversity alone, there is great variation across systems in the conditions under which infection and diversity are measured. In genetically homogenous bumble bee (*Bombus terrestris L.*) populations, *Nosema bombi* has higher success, but not *Crithidia bombi*, compared to diverse populations (19). In other cases, we see an increase in disease impact in homogenous host populations when infection is by multiple parasite species (19–22) but not always with specific interactions between one host-parasite species pair (23,24). Thirdly, because parasite infection is measured differently across studies, and even within systems, there is the potential that the relevant measure of parasite success isn’t used. For example, in honeybee (*Apis mellifera)* host populations, genetic diversity has a negative impact on parasite success when infection prevalence or parasite load is measured, but not always when host survival is calculated (25). Host survival might be less informative, particularly for parasites that are not obligate killers: not all hosts that are infected might die, but also host mortality can impede parasite transmission if the parasite requires host-to-host contact for infection to spread. It is therefore unclear whether the monoculture effect is relevant to host-parasite interactions across the tree of life.

We tested the generality of the monoculture effect with a formal meta-analysis across a range of host-parasite systems. We searched the published literature for all publicly available data sources and compared the effects of low and high host genetic diversity on parasite success using a nested random mixed effects meta-analysis model and Pearson’s correlation coefficient effect size *r* (with positive values indicating monoculture effects). We define ‘parasite success’ as a parasite’s ability to have a high abundance in the host population whether it is measured as infection load/host, prevalence, or host mortality. We also tested whether empirical contexts or biological factors associated with the species in the interaction could explain variation in the effect of diversity on parasite success.

## Materials and methods

### Literature search

Using Web of Knowledge, Google Scholar and PubMed, we searched the literature using various combinations of the following keywords: ‘host genetic diversity’, ‘low versus/and high host genetic diversity’, ‘heterogeneous versus/and homogenous host populations’, ‘monoculture effect’, ‘disease spread’, and ‘parasite prevalence’ to investigate the effect of low versus high host population diversity on parasite disease impact (see Supp. Fig. 1 for PRISMA flowchart (26) summarising study collection process). We gathered data where measurements were taken of parasite success in host populations of varying genetic diversity. These measurements included; parasite load, parasite virulence, parasite abundance, host mortality rate, viral concentrations, viral load, infection rate, and infection intensity. We also checked reference lists along with paper citations for other potential papers. Studies were also searched for and extracted from review papers.

Papers were included in this study if they met the following inclusion criteria:

i. The study was published in a peer reviewed academic journal.
ii. The study collected parasite success data from two distinct comparable host population groups with any measured difference in diversity, such as low versus high diversity, inbred versus outbred, and monoculture versus polyculture.
iii. In the study, both host population groups contained the same species.
iv. The study measured genetic diversity at the host population level and not community diversity or individual-level genetic heterozygosity.
v. The study was not conducted in an agricultural system.
vi. The study did not interfere with parasite or host lifecycle, as in passaging manipulations.

We decided to exclude agricultural studies as a meta-analysis has already demonstrated the benefits of intraspecific diversity to crop yields (and thus host fitness) in the presence of infectious disease (6).

### Statistical analysis

We calculated Pearson’s correlation coefficient, *r*, from studies using the method described in Field & Gillet (2010). This measure was chosen as it allowed for a direct comparison between two continuous variables, which in our case is low vs high host population diversity. To calculate effect size *r*, mean parasite infection measurements and their standard deviation for each treatment were extracted in the order of low host population diversity and high host population diversity. We extracted data from either paper figures, reported statistics in the text, or raw data received from authors. Where means and standard deviations in each group were not available (2 out of 22 studies), t-values and degrees of freedom were extracted.

We performed a nested random mixed effects meta-analysis model using the *rma.mv* function in the package *metafor* in R version 3.6.0 (R core development team). We chose this model to account for the fact that we collected several effect sizes per study, where some studies shared the same host species, which has the potential for pseudo-replication and phylogenetic non-independence. We first tested for an overall relationship between host population genetic diversity and parasite success using the entire dataset. Next, we tested for context dependence in the magnitude of the monoculture effects by focusing on the moderator variables: empirical environment, parasite infection measure, host reproduction, parasite functional group, host range, initial parasite diversity, and ability of parasite to cause host death. The measure of heterogeneity of moderator variables was reported as Q, where Q is the weighted sum of squares about the fixed effect estimate between subgroups (27).

We tested for an effect of empirical contexts or approach on the strength of the monoculture effect. In addition to dividing up studies into field or lab empirical environments, we also tested an effect of the parasite success measure on the strength of the monoculture effect. Thus, we separated measures into three groups; parasite prevalence, parasite load, and host mortality. Studies looking at overall parasite presence in a host population were placed under the category ‘parasite prevalence’. Where measures of parasite propagules per host were taken, studies were placed under ‘parasite load’. Measures of mortality within a population were placed under ‘mortality’. Measures of host survival were transformed into host mortality by subtracting calculated survival data from the entire measured population.

We then focused on the impact of aspects of host and parasite biology that could explain variation in the effect of host diversity on parasite success. Specifically, we tested whether the strength of the monoculture effect was related to host reproductive mode, given sexual and asexual strategies generate disparate levels of genetic diversity; infection by micro- or macroparasites, as the former tends to be associated with higher pathogenicity (28); and finally, host range (specialists or generalists), as it is assumed host resistance is genetic-based and there is a long-standing association between host and parasite. Here, we define specialist as a parasite only able to infect one host species and generalist as a parasite able to infect multiple host species. In addition, because higher levels of parasite diversity are thought to increase the pool of susceptible hosts in a diverse population, we separated studies into three categories – one genotype of one parasite species (1 Genotype), multiple parasite genotypes of one parasite species (>1 Genotypes), and many parasite species (>1 Species) – to determine the importance of parasite diversity on the strength of the monoculture effect. Lastly, we tested whether the parasite’s ability to cause host death was associated with the strength of the monoculture effect. More virulent parasites could select for greater levels of resistance in the host population, whereas there may not be genetic variation for resistance in diverse host populations infected by less harmful parasites.

### Assessing for potential publication bias

Studies that report larger effects are more likely to get published in comparison to studies reporting smaller effects (29). To check for publication bias, we visualised the spread of our effect sizes by creating a funnel plot (Supp. Fig. 2). We then performed a Fail-Safe N analysis to calculate the number of additional studies needed to reduce the significance level of the weighted average effect size (30).

## Results

We found 22 papers containing data to answer the research question and followed the inclusion criteria. Papers often included results from multiple experiments or exposures to multiple parasite species. A total of 66 effect sizes were retrieved from this data set, covering a diverse range of host and parasite species (Table 1).

**Table 1:**
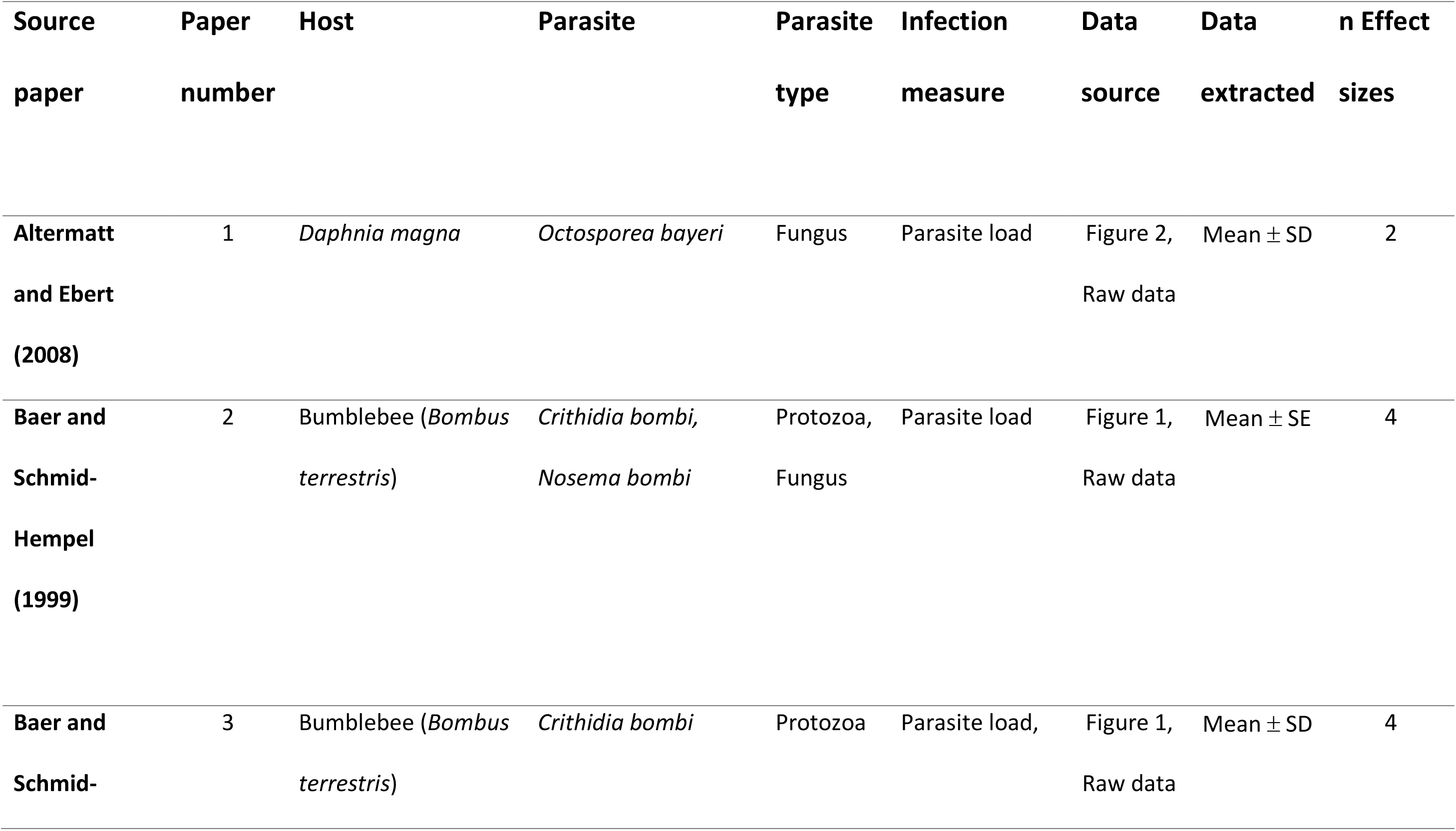
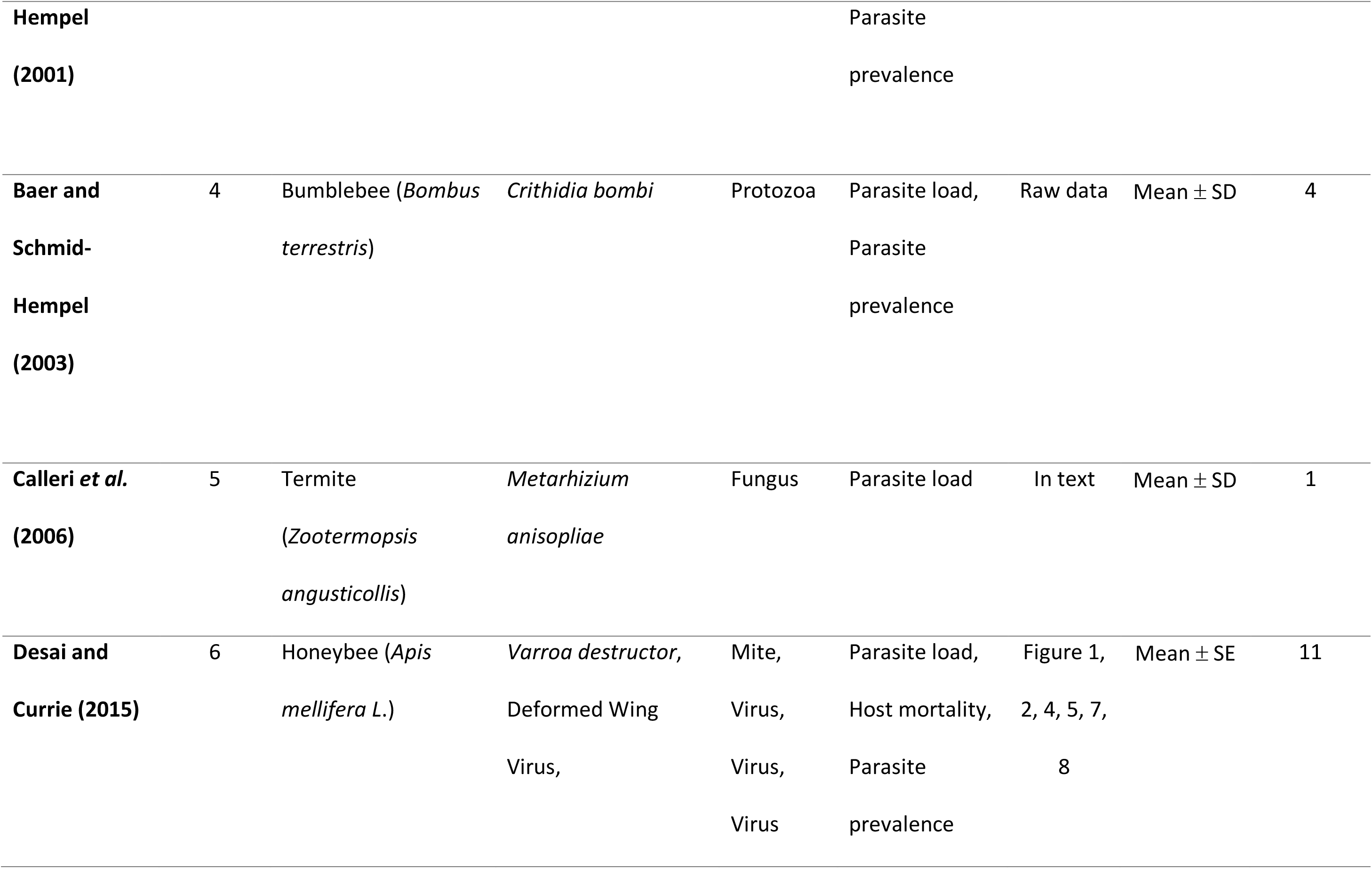
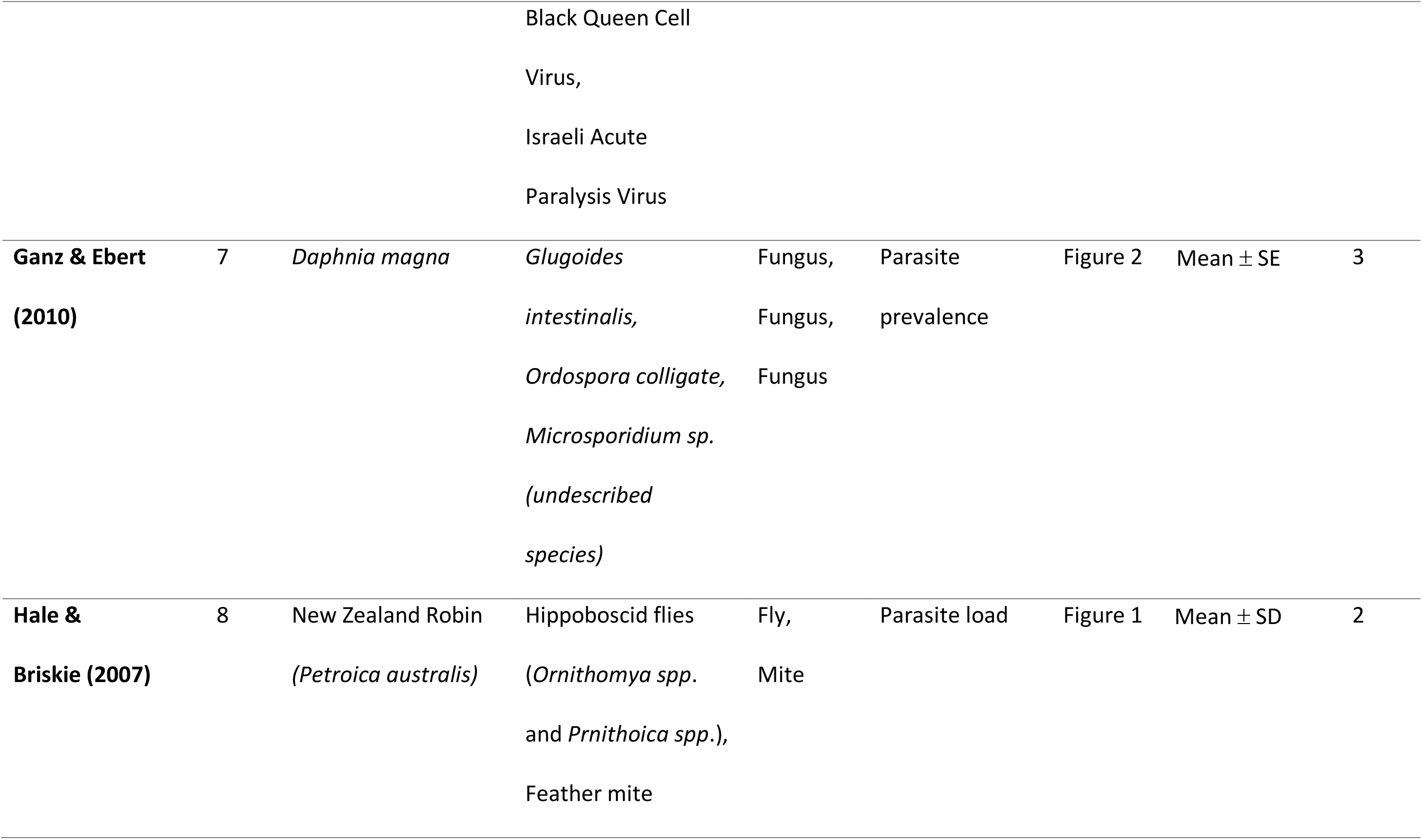
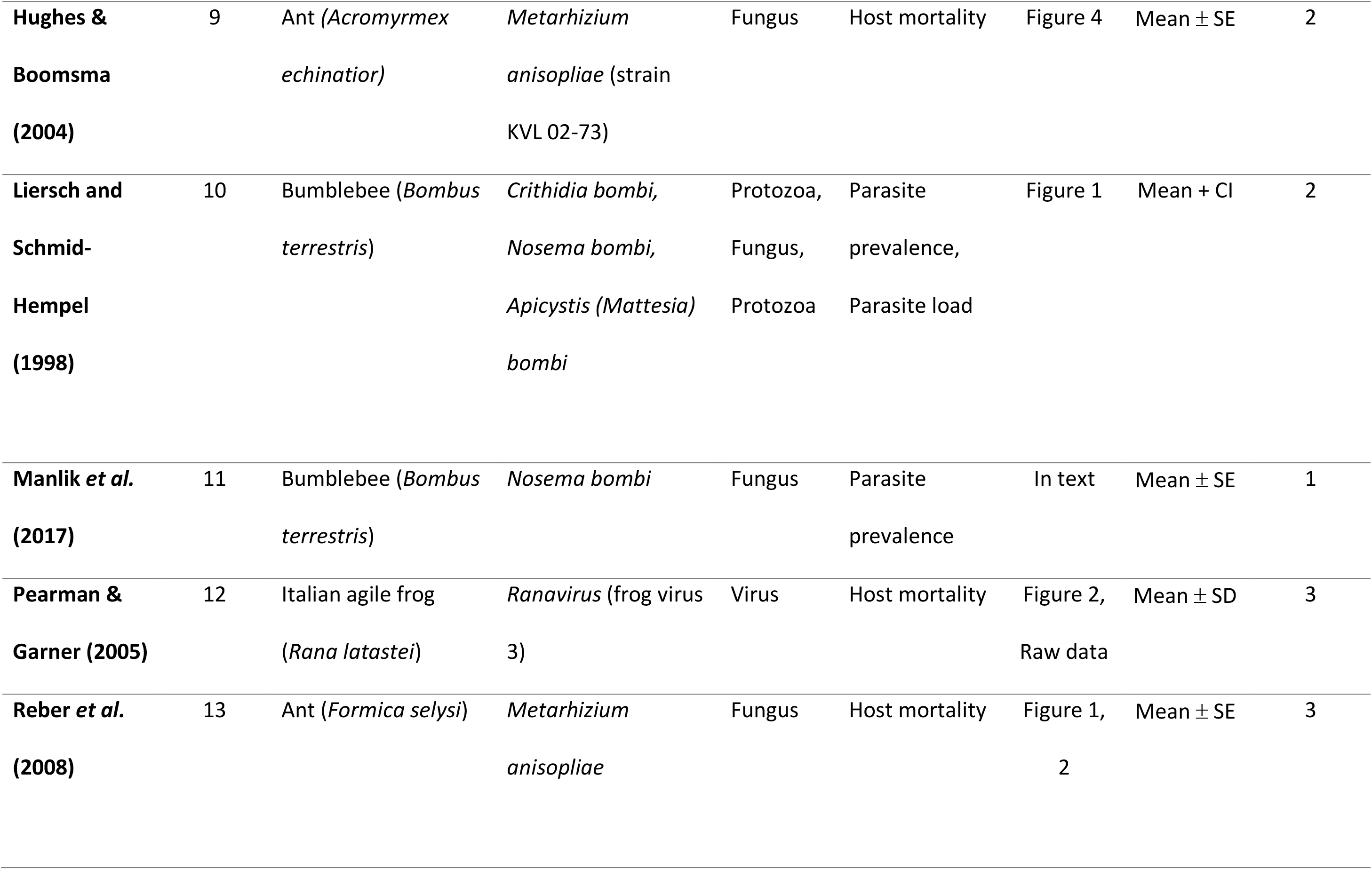
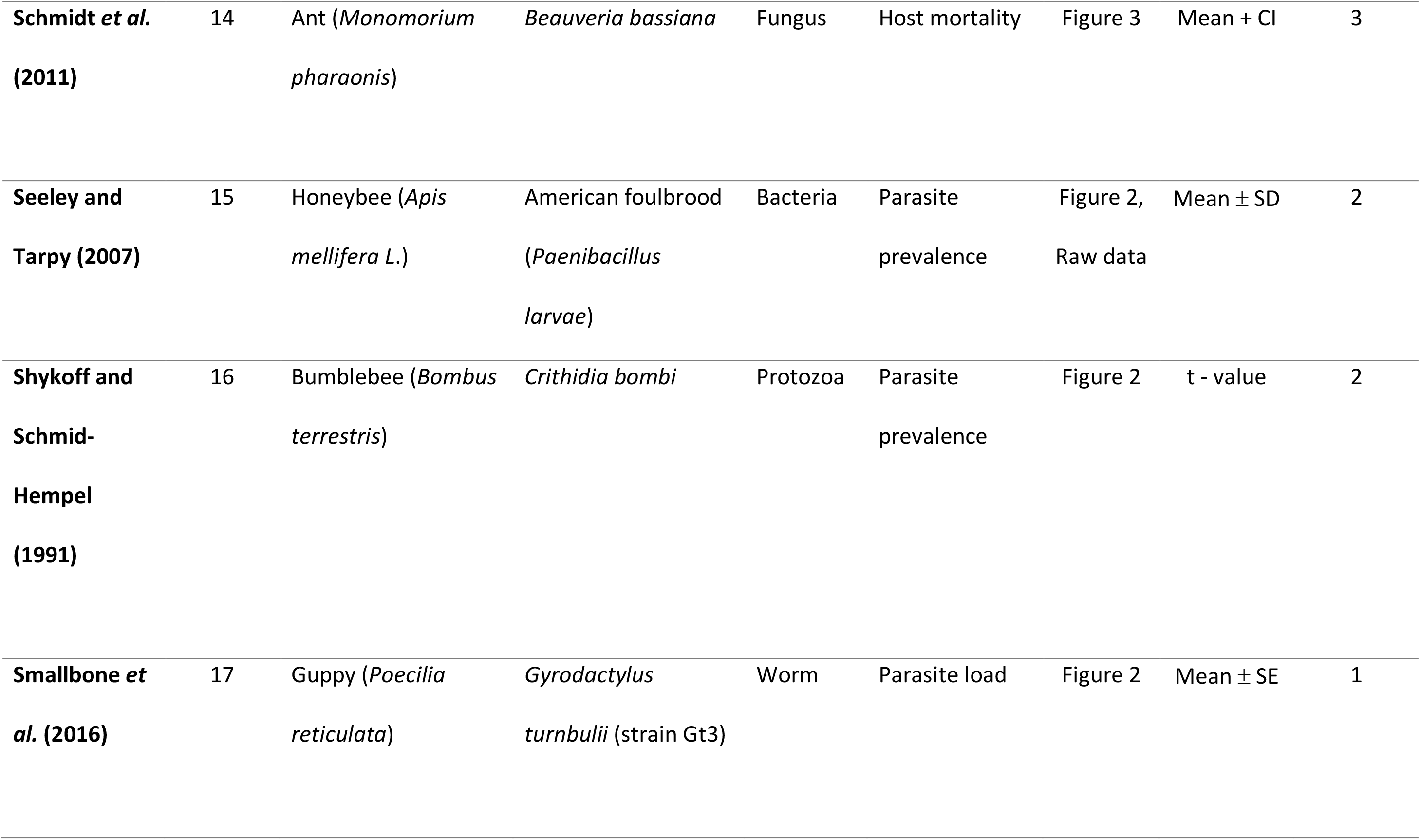
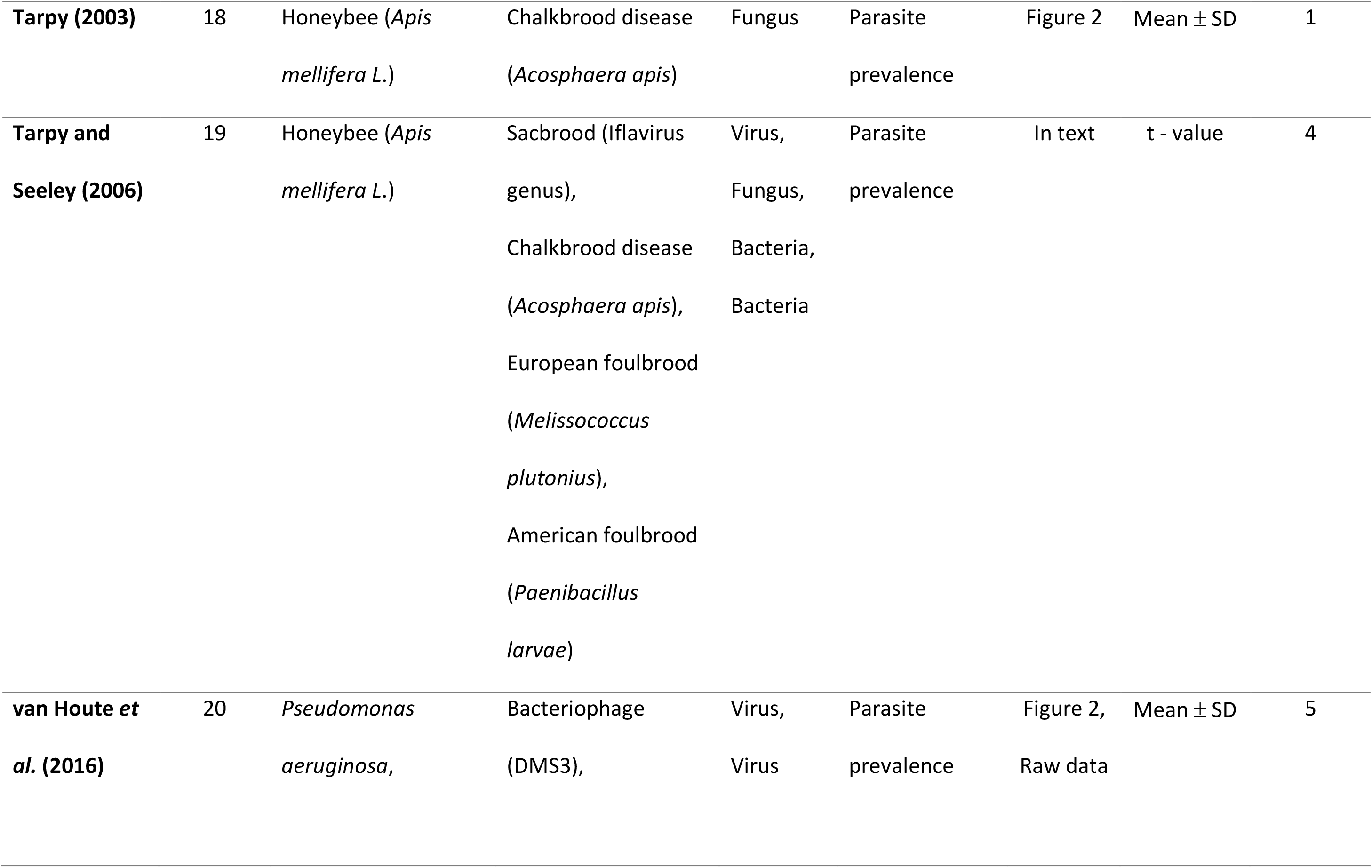
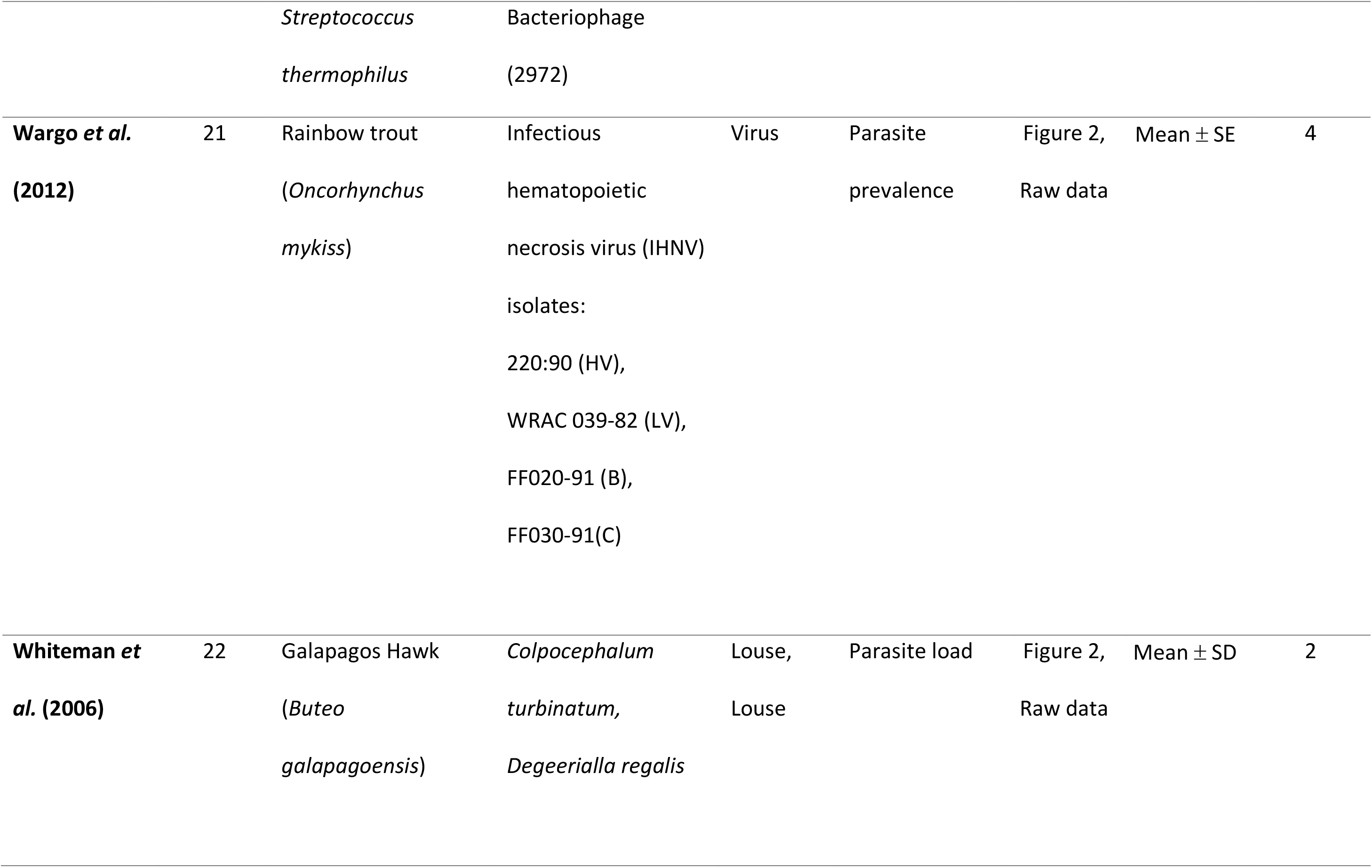
Summary of literature on the effect of host population genetic diversity on measures of parasite success across host-parasite systems.

After the construction of a funnel plot, we find no indication of a publication bias in this meta-analysis data set, with the majority of points falling within the plot (Supp. Fig. 1). Rosenberg’s Fail-safe N analysis showed that an additional 644 studies would need to be added to reduce the significance level of this meta-analysis.

Our results are consistent with the monoculture effect hypothesis, showing that low host genetic diversity increases parasite success (r = 0.3950, z = 3.1349, p < 0.0001, Fig. 1A). We found that the strength of the direction of the effect size is influenced by empirical environment (Q = 8.4778, d.f. = 1, p = 0.0036, Fig. 1B), where field studies (r = 0.2801) did significantly differ from lab studies (r = 0.1077). However, parasite infection measures (i.e. parasite load, parasite prevalence, or host mortality) do not significantly influence the effect size (Q = 3.5302, d.f. = 2, p = 0.1712, Fig. 1C).

**Figure 1:**
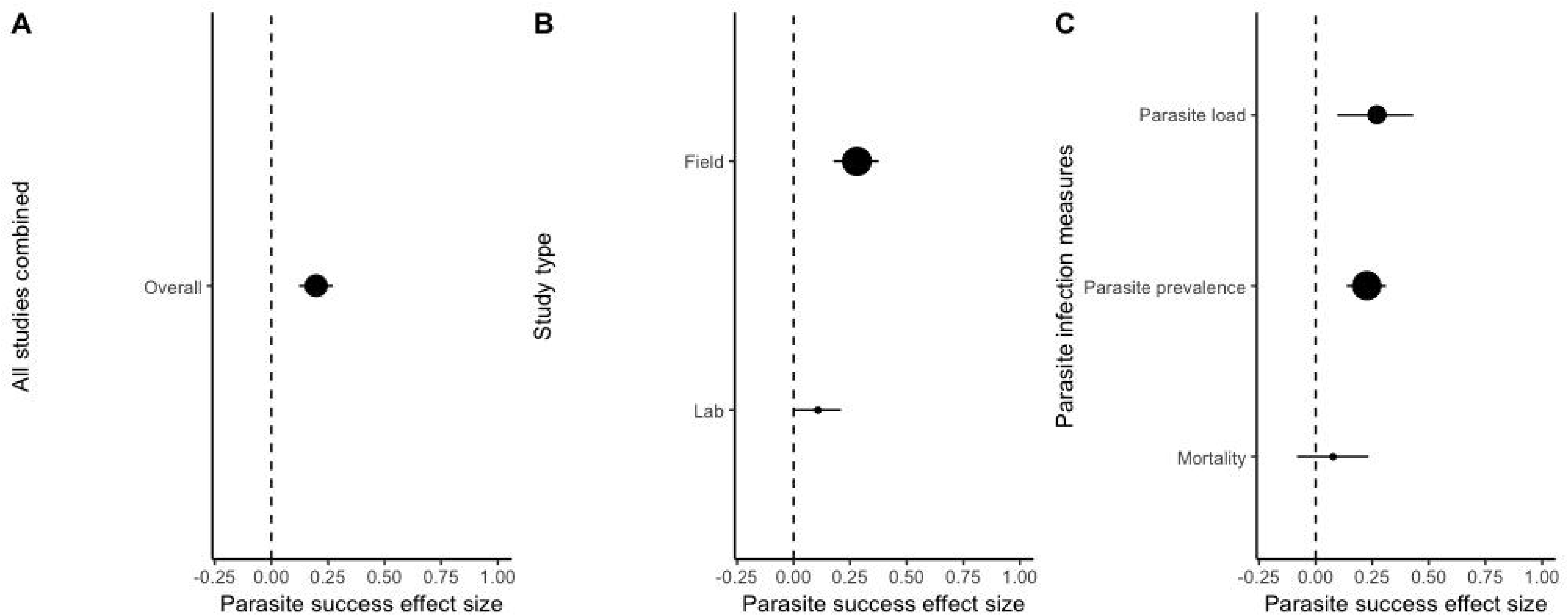
Impact of study approach on the effect of host genetic diversity on parasite success. Positive values indicate a monoculture effect is present (i.e., a negative association between genetic diversity and parasite success). Negative values represent the opposite relationship. At an effect size of zero (dashed line), there is no relationship between host genetic diversity and parasite success. (A) Overall effect size (n = 66). (B) Moderator analysis of study type between field (n = 36) and lab (n = 30) studies. (C) Moderator analysis of parasite infection measures between parasite load (n = 19), parasite prevalence (n = 34), and host mortality (n = 13). The size of the dot corresponds to the sample size. Effect sizes are shown with 95% confidence intervals.

We examined the impact of a suite of host and parasite characteristics on the strength of the monoculture effect. We found that host reproduction was not a factor that significantly influenced the strength of the effect size (Q = 3.7744, d.f. = 2, p = 0.1515, Fig. 2A). A study by Altermatt & Ebert (2008) followed parasite infection of *Daphnia* during both sexual and asexual reproduction, and was thus placed as a separate variable. We then focused on parasite characteristics, we found that parasite functional group significantly influenced the strength of the direction of the effect size (Q = 8.7057, d.f. = 1, p = 0.0032, Fig. 2B). Where macroparasites (r = - 0.0091) had mostly no or a slightly negative impact, but microparasites (r = 0.2298) showed a strong, positive impact. The direction of the effect size was found not to be influenced by host range (Q = 0.2771, d.f. = 1, p = 0.5986, Fig. 2C). We also found that parasite diversity was not a significant factor on the strength of the monoculture effect (Q = 3.5302, d.f. = 2, p = 0.1712, Fig. 2D). Finally, we investigated whether the ability of a parasite to cause host mortality would influence the direction of the effect size. We found a significant effect on parasite success (Q = 3.8744, d.f. = 1, p = 0.0490, Fig. 2E), whereby studies using parasites that could kill hosts showed a stronger monoculture effect (r = 0.2120) than those with less virulent parasites (r = 0.0627).

**Figure 2:**
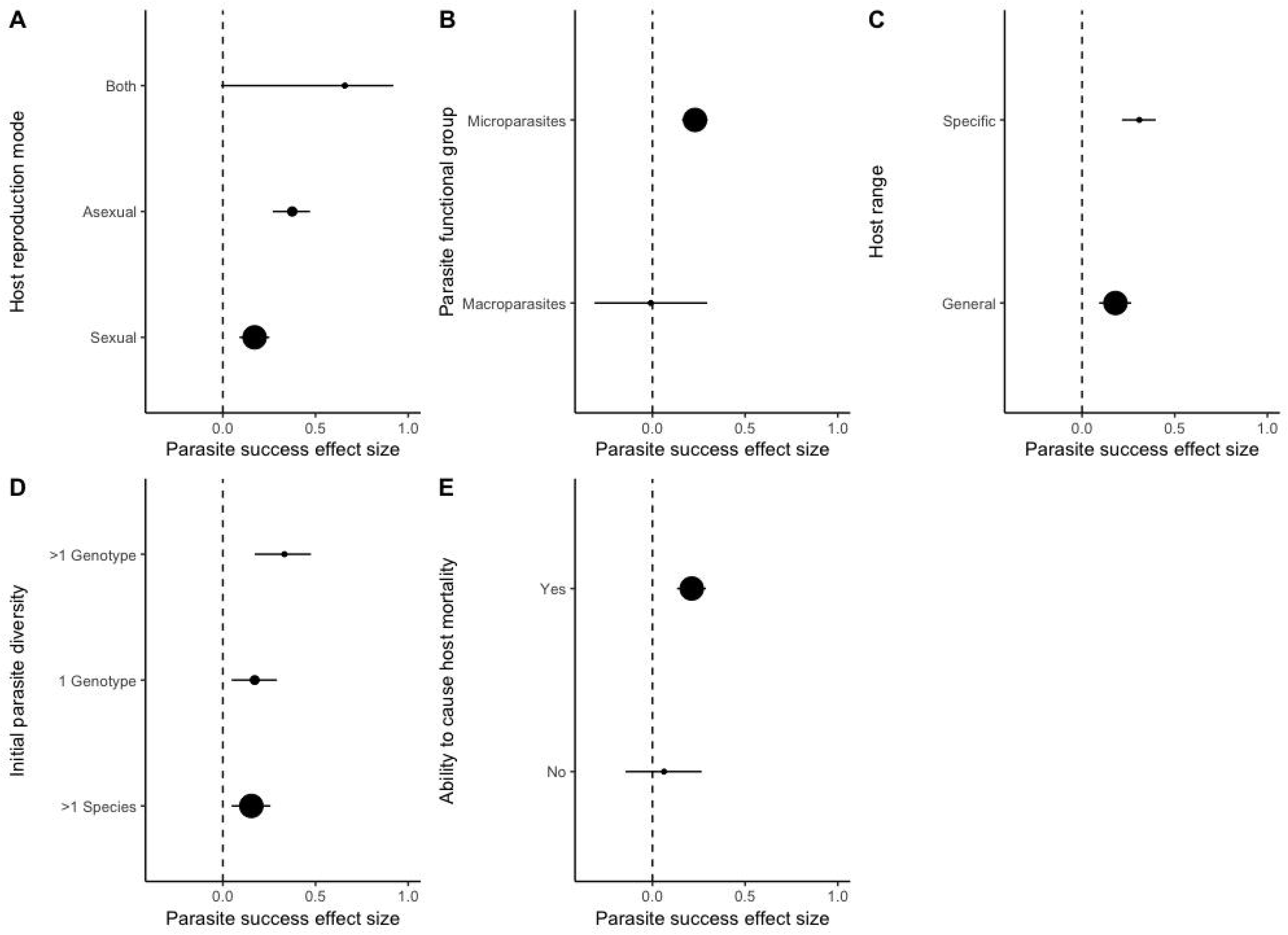
Impact of host and parasite characteristics on the effect of host genetic diversity on parasite success. Positive values indicate a monoculture effect (i.e., a negative association between genetic diversity and parasite success). Negative values represent the opposite relationship. The dashed line (effect size of zero) represents no relationship between host genetic diversity and disease spread. Moderator analysis of (A) host reproduction mode: asexual (n = 5), both (n = 2), and sexual (n = 59) effect sizes, (B) of parasite functional group between microparasite (n = 56) and macroparasite (n = 10) effect sizes, (C) host range between specific (n = 15) and general (n = 51) parasite effect sizes, (D) initial parasite diversity between >1 genotype (n = 14), 1 genotype(n = 15), and >1 species (n = 37) effect sizes, and (E) of the ability of a parasite to cause host death, displayed as yes (n = 56) and no (n = 10) effect sizes. The size of the dot corresponds to the sample size. Effect sizes are shown with 95% confidence intervals.

## Discussion

Our meta-analysis shows that host population genetic diversity reduces parasite success across multiple systems, approaches, and environments. Indeed, the monoculture effect is revealed under the majority of the biological variables we tested in the host-parasite relationship, but that microparasites and parasites that kill are more likely to encounter differences in resistance in host populations varying in diversity. Our findings additionally highlight the potential damage that emerging infectious diseases may have on genetically homogenous host populations, given that the monoculture effect is not dependent on a parasite’s host range.

The parasites included in our meta-analysis were highly variable in terms of their host range. However, we show that the monoculture effect is independent of a parasite’s host range. Indeed, the monoculture effect is equally as prevalent in highly specialised interactions (31–33), in broad spectrum interactions at the genotypic level (34), and in those that cross host-species boundaries (21,22,35). That host range is not a factor here is in contrast to those results found in crop studies. For example, in rusts and powdery mildews, disease severity is driven by pathogen specificity (5). The mirroring of parasite virulence genes to host resistance genes means that crop mixtures need to contain both susceptible and resistant cultivars to avoid a monoculture effect. When there is a lack of host specificity, mixed cultivar populations are just as susceptible as monocultures. For example, mixed cultivar populations have been observed to be slightly more susceptible to infection (36) or completely susceptible (37) in comparison to monocultures to the fungal pathogen *Mycosphaerella graminicola*. These findings suggest that the threat to crops from generalist parasites is greater than specialist parasites.

Given that host range did not influence the strength of the monoculture effect, it is possible that novel parasites, just as adapted parasites, could have high success in host monocultures. Essentially, homogenous populations could be vulnerable to outbreaks with spill-over or emerging infectious diseases which are less likely to be host specific (38), but for which there is clearly genetic variation for resistance. The resistance to emerging parasites in these cases could be due to historical contact or similar mechanisms of infection to parasites with an evolutionary history with the host (39). Nevertheless, this result is concerning from a conservation perspective as global climate change has the potential to reduce within-species genetic diversity (40) and alter host population ranges (41,42). Natural movement of individuals between populations has always served to bolster host diversity (42), and introducing new genotypes is an approach applied by conservation biologists to improve population viability (10). Whilst adding individuals to a population could increase diversity and reduce inbreeding (43), a risk may be that new individuals bring in new parasites to the population (44). Given that we found a stronger effect in field studies, these consequences are of real concern. The potential being an increased overlap between host populations with low genetic diversity and novel infections.

The fact that we found a stronger monoculture effect in field studies highlights the importance of the maintenance of diversity in natural populations. As hosts are exposed to a greater variety of parasites in the field, there could be higher levels of resistance already present in diverse populations (39). Thus, when host diversity is artificially reduced (21), parasites normally unable to rapidly spread through a host population can now infect with minimal selection on virulence evolution. In addition, secluded host populations, such as island populations of Galapagos hawks (22), are naturally considered inbred compared to their main land or larger island counter parts and are therefore more vulnerable to infection. Also, island populations as well as social insects, such as bees (45), ants (46), and termites (47), live in tight proximities to each other making parasite transmission easier in homogenous populations. Indeed, despite being subjected to environmental noises, the monoculture effect is strong in the natural environment.

In our meta-analysis, macroparasites were not impeded by genetic heterogeneity in host populations. The macroparasites in the studies included herein are all ectoparasites, and their biology may explain why. Their transmission is often dependent on host-to-host contact (48,49) and thus host density is a critical factor in parasite success (48). Host density may play a more important role than host genetic diversity such that similarly aggregated populations of either genetically high or low host populations might be equally susceptible to infection. It has been shown that clustering of captive animal populations restricted by movement or wild animal populations restricted by ranges are highly vulnerable to ectoparasites (44,50). Moreover, host social behaviours, such as grooming (25) or preening (22) can reduce ectoparasite success. In fact, in populations where social grooming is correlated with relatedness, ectoparasite load is dramatically reduced in highly related individuals (51). Taken together, host diversity on its own does not always explain a reduction in parasite success, particularly in the case of ectoparasites.

We reveal that the monoculture effect is more likely to be observed in systems with a parasite that can cause host mortality. This outcome may stem from greater selection for resistance in diverse host populations at risk of infection and death from parasites (52). Whilst some parasites in the relevant studies are obligate killers, such as bacteriophages (33), some merely have the potential to cause host mortality. For example, *Crithidia bombi* can cause in mortality in bumble bees (*Bombus* spp) when the colony is stressed by lack of access to food sources (53). It is nevertheless possible that host population genetic diversity, as measured in the studies with less virulent parasites, may not be correlated with diversity in resistance *per se*.

Understanding the impact of reduced genetic diversity on parasite infection outside of agricultural systems is crucial because of anthropogenic threats to the diversity of wild populations. This meta-analysis reveals that the monoculture effect is a widespread phenomenon across host and parasite species in nature, with microparasites and host-killing parasites being the most likely to encounter resistance in diverse host populations. Indeed, these broad patterns show that genetic diversity is a robust weapon against infection, but that further attacks on diversity could drive outbreaks of both coevolving and emerging infectious diseases. However, these results suggest that conservation efforts should focus on preserving population genetic diversity in vulnerable populations to improve their ability to fight off deadly infections.

## Acknowledgments

We are grateful to PB Pearman, AR Wargo, P Schmid-Hempel, NK Whiteman, TD Seeley, DR Tarpy, F Altermatt, S van Houte & E Westra for sharing their raw data with us. We also thank CM Lively for comments on our manuscript. A.K.E.E. acknowledges funding from Natural Environment Research Council (NE/L002612/1). Funding was also provided by the Leverhulme Trust and European Research Council to C.R-M. and K.C.K.

