## Supplemental Figure 1 for "Diversity and disease: evidence for the monoculture effect beyond agricultural systems"

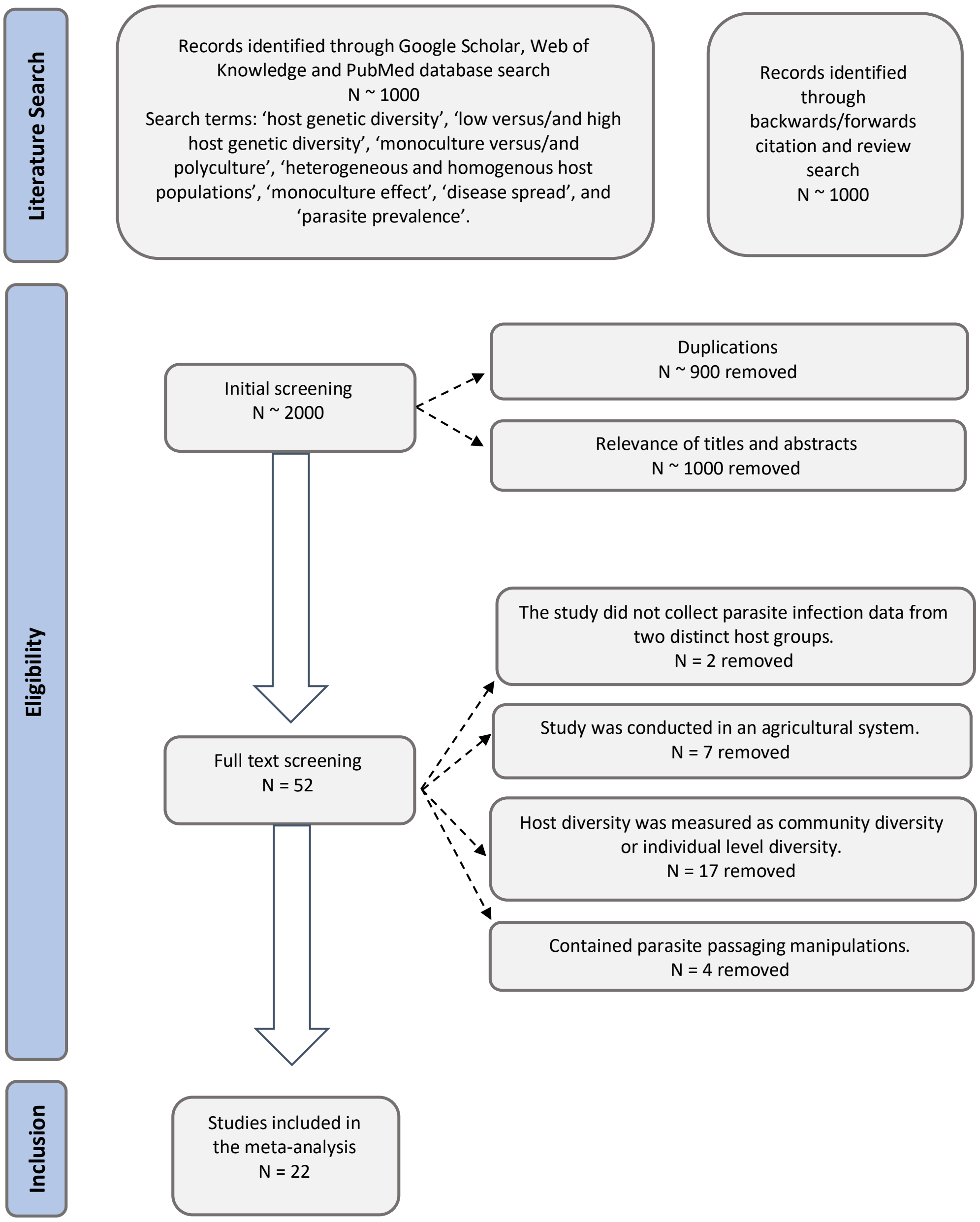


**Supplementary Figure 1:** PRISMA flow chart of literature search and study selection process. Studies excluded from the analysis are listed in Supplementary Table 2.
