## Supplemental Figure 2 for "Diversity and disease: evidence for the monoculture effect beyond agricultural systems"

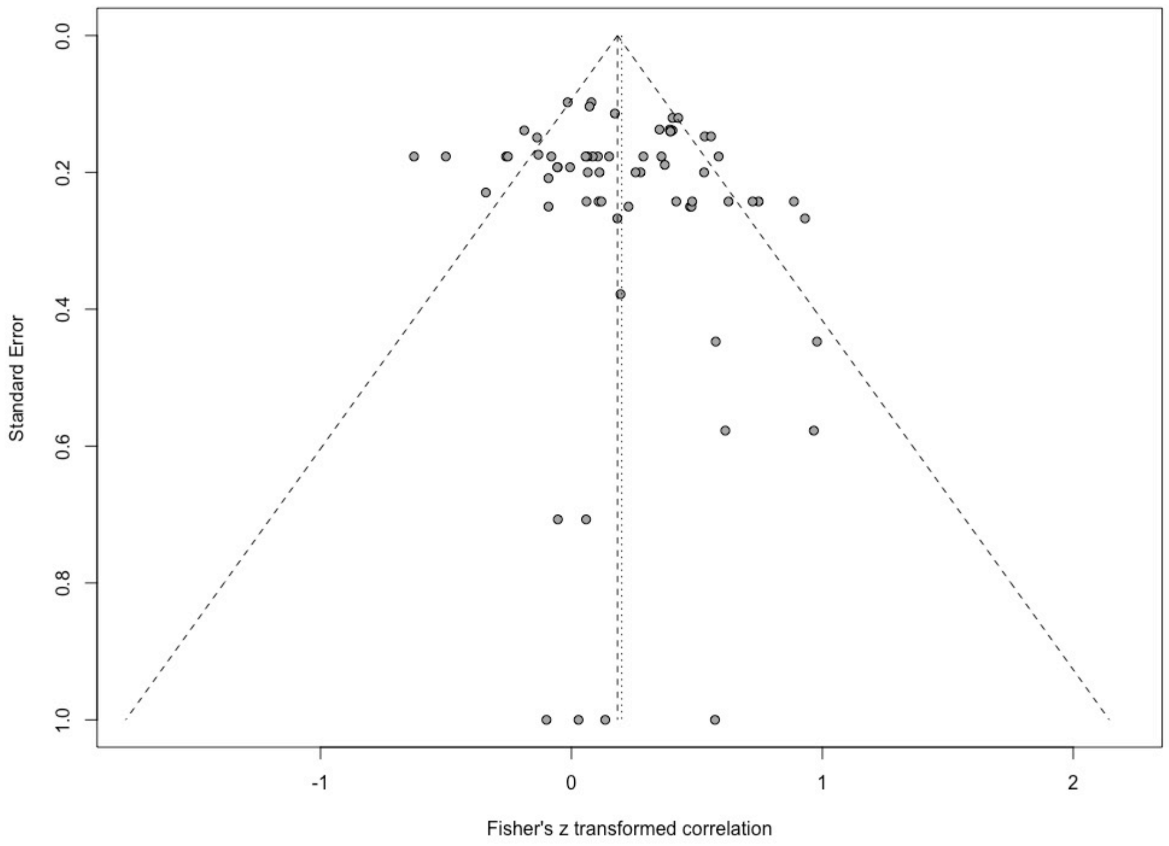


**Supplementary Figure 2:** Funnel plot of the meta-analysis data set. Points on the graph represent the relationship between host genetic diversity and disease impact for each study. A vertical line has been added to show the population effect size estimates, and the diagonal lines show the 95% confidence interval of this estimate.
