## Supplemental Table 1 for "Diversity and disease: evidence for the monoculture effect beyond agricultural systems"

**Supplementary Table 1:** Studies excluded from meta-analysis

| Study | Reason for exclusion |
|---|---|
| **Abdala-Roberts**, L., Gonzalez-Moreno, A., Mooney, K.A., Moreira, X., Gonzalez-Hernandez, A. & Parra-Tabla, V. (2016) Effects of tree species diversity and genotypic diversity on leafminers and parasitoids in a tropical forest plantation. Agricultural and Forest Entomology, 18, 43-51. | Host diversity was measured as community diversity. |
| **Acevedo-Whitehouse**, K., Gulland, F., Greig, D. & Amos, W. (2003) Disease susceptibility in California sea lions. Nature, 422, 35. | Host diversity was measured as individual level diversity. |
| **Agha**, R., Gross, A., Rohrlack, T. & Wolinska, J. (2018) Adaptation of a Chytrid Parasite to Its Cyanobacterial Host Is Hampered by Host Intraspecific Diversity. Frontiers in Microbiology, 9:921. | Contained parasite passaging manipulations. |
| **Alexander**, H.M., Roelfs, A.P. & Cobbs, G. (1986) Effects of disease and plant competition on yield in monocultures and mixtures of two wheat cultivars. Plant Pathology, 35, 457-465. | Study was conducted in an agricultural system. |
| **Andras**, J.P. (2017) Genetic variation of the Caribbean sea fan coral, *Gorgonia ventalina*, correlates with survival of a fungal epizootic. Marine Biology, 164:130. | Host diversity was measured as individual level diversity. |
| **Browning**, J.A. & Frey, K.J. (1969) Multiline cultivars as a means of disease control. Annual Review of Phytopathology, 7, 355-382. | Study was conducted in an agricultural system. |
| **Campbell**, G., Noble, L.R., Rollinson, D., Southgate, V.R., Webster, J.P. & Jones, C.S. (2010) Low genetic diversity in a snail intermediate host (*Biomphalaria pfeifferi* Krass, 1848) and schistosomiasis transmission in the Senegal River Basin. Molecular Ecology, 19, 241-256. | Host diversity was measured as individual level diversity. |

| | |
|---|---|
| **Ellison**, A., Cable, J. & Consuegra, S. (2011) Best of both worlds? Association between outcrossing and parasite loads in a selfing fish. Evolution, 65, 3021-3026. | Host diversity was measured as individual level diversity. |
| **Hendrick**, P.W., Kim, T.J & Parker, K.M. (2001) Parasite resistance and genetic variation in the endangered *Gila topminnow*. Animal Conservation Forum, 4, 103-109. | Host diversity was measured as individual level diversity. |
| **Hughes**, W.O.H. & Boomsma, J.J. (2006) Does genetic diversity hinder parasite evolution in social insect colonies? Journal of Evolution Biology, 19, 132-143. | Contained parasite passaging manipulations. |
| **Jinks**, J.L. & Grindle, M. (1963) Changes induced by training in *Phytophthora infestans*. Heredity, 18, 245-264. | Contained parasite passaging manipulations. |
| **Kerstes**, N.A.G. & Wegner, K.M. (2011) The effect of inbreeding and outcrossing of Tribolium castaneum on resistance to the parasite *Nosema whitei*. Evolutionary Ecology Research, 13, 681-696. | Host diversity was measured as individual level diversity. |
| **Knott**, E.A. & Mundt, C.C. (1990) Mixing ability analysis of wheat cultivar mixtures under diseased and nondiseased conditions. Theoretical and Applied Genetics, 80, 313-320. | Study was conducted in an agricultural system. |
| **Kubinak**, J.L., Cornwall, D.H., Hasenkrug, K.J., Adler, F.R. & Potts, W.K. (2014) Serial infection of diverse host (Mus) genotypes rapidly impedes pathogen fitness and virulence. Proceedings of the Royal Society B: Biological Sciences, 282: 20141568. | Contained parasite passaging manipulations. |
| **Lai**, R., You, M., Zhu, C., Gu, G., Lin, Z., Liao, L., Lin, L. & Zhong, X. (2017) Myzus persicae and aphid-transmitted viral disease control via variety intercropping in flue-cured tobacco. Crop protection, 100, 157-162. | Study was conducted in an agricultural system. |
| **Liao**, H., Luo, W., Pal, R., Peng, S. & Callaway, R.M. (2016) Context-dependency | |

| | |
|---|---|
| and the effects of species diversity on ecosystem function. Biological Invasions, 18, 3063-3079. | Host diversity was measured as community diversity. |
| **Lopez-Uribe**, M.M., Appler, R.H., Youngsteadt, E., Dunn, R.R., Frank, S.D. & Tarpy, D.R. (2017) Higher immunocompetence is associated with higher genetic diversity in feral honey bee colonies (*Apis mellifera*). Conservation Genetics, 18, 659-666. | Host diversity was measured as individual level diversity. |
| **Meagher**, S. (1999) Genetic diversity and *Capillaria hepatica* (Nematoda) prevalence in Michigan deer mouse populations. Evolution, 53, 1318-1324. | Host diversity was measured as individual level diversity. |
| **Meyer-Lucht**, Y., Otten, C., Puttker, T., Pardini, R., Metzger, J.P.& Sommer, S. (2010) Variety matters: adaptive genetic diversity and parasite load in two mouse opossums from the Brazilian Atlantic forest. Conservation Genetics, 11, 2001-2013. | Host diversity was measured as individual level diversity. |
| **Mitchell**, C.E., Tilman, D. & Groth, J.V. (2002) Effects of grassland plant species diversity, abundance, and composition on foliar fungal disease. Ecological Society of America, 83, 1713-1726. | Host diversity was measured as community diversity. |
| **O'Brien**, S.J., Roelke, M.E., Marker, L., Newman, A., Winkler, C.A., Meltzer, D., Colly, L., Evermann, J.F., Bush, M. & Wildt, D.E. (1985) Genetic basis for species vulnerability in the cheetah. Science, 227, 1428-1434. | Host diversity was measured as individual level diversity. |
| **Palmer**, K.A. & Oldroyd, B.P. (2003) Evidence for intra-colonial genetic variance in resistance to American foulbrood of honey bees (*Apis mellifera*): further support for the parasite/pathogen hypothesis for the evolution of polyandry. Naturwissenschaften, 90, 265-268. | Host diversity was measured as individual level diversity. |
| **Pilet**, F., Chacon, G., Forbes, G.A., Andrivon, D. (2006) Protection of susceptible potato | |

| | |
|---|---|
| cultivars against late blight in mixtures increases with decreasing disease pressure. Phytopathology, 96, 777-783. | Study was conducted in an agricultural system. |
| **Rottstock**, T., Joshi, J., Kummer, V. & Fischer, M. (2014) Higher plant diversity promotes higher diversity of fungal pathogens, while it decreases pathogen infection per plant. Ecological Society of America, 95, 1907-1917. | Host diversity was measured as community diversity. |
| **Schmid-Hempel**, P. & Crozier, R.H. (1999) Polygyny versus polyandry versus parasites. Philosophical Transactions of the Royal Society Biological Sciences, 354, 507-515. | Host diversity was measured as individual level diversity. |
| **Spielman**, D., Brook, B.W., Briscoe, D.A. & Frankham, R. (2004) Does inbreeding and loss of genetic diversity decrease disease resistance? Conservation Genetics, 5, 439-448. | Host diversity was measured as individual level diversity. |
| **Velo-Anton**, G., Rodriguez, D., Savage, A.E., Parra-Olea, G., Lips, K.R. & Zamudio, K.R. (2012) Amphibian-killing fungus loses genetic diversity as it spread across the New World. Biological Conservation, 146, 213-218. | The study did not collect parasite infection data from two distinct host groups. |
| **Whitehorn**, P.R., Tinsley, M.C., Brown, M.J.F., Darvill, B. & Goulson, D. (2011) Genetic diversity, parasite prevalence and immunity in wild bumblebees. Proceedings of the Royal Society B: Biological Sciences, 278, 1195-1202. | The study did not collect parasite infection data from two distinct host groups. |
| **Yang**, B., Ge, F., Ouyang, F. & Parajulee, M. (2012) Intra-species Mixture Alters Pest and Disease Severity in Cotton. Environmental Entomology, 41, 1029-1036. | Study was conducted in an agricultural system. |
| **Zhu,** Y., Chen, H., Fan, J., Wang, Y., Li, Y., *et al*. (2000) Genetic diversity and disease control in rice. Nature, 406, 718-722. | Study was conducted in an agricultural system. |
